# Fluctuating selection strength and intense male competition underlie variation and exaggeration of a water strider’s male weapon

**DOI:** 10.1101/430314

**Authors:** William Toubiana, Abderrahman Khila

## Abstract

Sexually selected traits can reach high degrees of expression and variation under directional selection. A growing number of studies suggest that such selection can vary in space, time and form within and between populations. However, the impact of these fluctuations on sexual trait expression is poorly understood. The water strider *Microvelia longipes* displays a striking case of exaggeration and phenotypic variation where males display extreme differences in the size of their rear legs. To study the origin and maintenance of this exaggerated trait, we conducted comparative behavioral and morphometric experiments in a sample of *Microvelia* species. We uncovered differences both in the mating behavior and the degree of sexual dimorphism across these species. Interestingly, *M. longipes* evolved a specific mating behavior where males compete for egg-laying sites, consisting of small floating objects, to intercept and copulate with gravid females. Field observations revealed rapid fluctuation in *M. longipes* habitat stability and the abundance of egg-laying sites. Through male-male competition assays, we demonstrated that male rear legs are used as weapons to dominate egg-laying sites and that intense competition is associated with the evolution of rear leg length exaggeration. Paternity tests using genetic markers demonstrated that small males could only fertilize about 5% of the eggs when egg-laying sites are limiting, whereas this proportion increased to about 20% when egg-laying sites become abundant. Furthermore, diet manipulation and artificial selection experiments also showed that the exaggerated leg length in *M. longipes* males is influenced by both genetic and nutritional factors. Collectively, our results highlight how fluctuation in the strength of directional sexual selection, through changes in the intensity of male competition, can drive the exaggeration and phenotypic variation in this weapon trait.

## Introduction

Phenotypic variation is central to the process of evolution [1], and understanding the mechanisms of its emergence and persistence in natural populations remains at the forefront of evolutionary biology studies [2], Sexually selected traits represent some of the primary examples illustrating both intra- and interspecies phenotypic variation [3, 4], Males in both vertebrates and invertebrates are known to wield extravagant phenotypes that can differ in their nature, location, size, and shape [3-5]. Examples include deer antlers, beetle horns, eyestalks in some flies, pseudoscorpion antennae and harlequin beetle legs. Some males of these species can develop degrees of trait expression so high that they appear exaggerated compared to other body parts or other homologous structures in the other sex [6]. A central prediction for these exaggerated traits to evolve is that only large individuals can afford to bear them, which can be a good indicator of body size and thus act as an honest signal for male quality [7–9]. Under this prediction, females will favor males with the highest trait expression, thus imposing strong directional selection in favor of trait exaggeration [3, 10]. In other situations, the trait is used as a weapon in male-male competition with its size being a good predictor for the outcome of the contest over access to females [11–14].

In these examples, sexual selection is thought to be directional and persistent over time [9, 12, 15]. These traits are also known to be subject to survivorship costs, which constrain their degree of expression resulting in a net stabilizing selection. These observations raise important questions regarding the maintenance of phenotypic variation in natural populations [3, 9, 15–21], A growing number of studies suggest that selection may not be as consistent over time and space, and that environmental changes may influence the strength, direction, and form of sexual selection [22–26], These fluctuations in selection may, in turn, favor the elevated plastic response and genetic variation observed in sexual traits, possibly influencing their variation and evolution over time [9, 21–23], Studies assessing the interplay between selection, genetics and plasticity, within the context of a changing environment are therefore crucial to the general understanding of the origin and maintenance of highly variable exaggerated sexual traits.

Here we focus on a novel model system, the water strider *Microvelia longipes*, that displays a strong sexual dimorphism where males have evolved both longer and more variable rear legs than females [27], The genus *Microvelia* (Heteroptera, Gerromorpha, Veliidae) comprises some 170 species of small water striders distributed worldwide and occupying various fresh water habitats including temporary rain puddles and stable large water bodies [27], First we reconstructed phylogenetic relationships of five *Microvelia* species and compared their degree of dimorphism, scaling relationships between leg and body length, and various aspects of mating behavior. We report a clear association between the intensity of male competition and the evolution of trait exaggeration in *M. longipes* males. We then determined the fitness advantages of these exaggerated legs through fertilization success performed under selective conditions reflecting fluctuations in their natural environment. Finally, we assessed the contribution of the strength of sexual selection, genetic variation, and phenotypic plasticity to the variation of exaggerated rear legs in *M. longipes* males.

## Results and discussion

### Sexual Dimorphism and scaling relationships in *Microvelia* species

We found a considerable inter-species variation in the degree of sexual dimorphism within the *Microvelia* genus (Figure 1). Measurements of various body parts revealed dimorphism in average body length, leg length, and the scaling relationship between these two traits (Figure 1B; Supplementary table 1). In some species, such as *M. americana* and *M. paludicola*, the dimorphism in leg and body length is small, whereas in others such as *M. longipes*, the dimorphism is spectacular (Figure 1A). The extreme leg elongation found in *M. longipes* males originates from the evolution of hyperallometry where the allometric coefficient is significantly higher than 1 and reaches a value of 3.2 - one of the highest known (Figure 1B; Supplementary table 1) [5, 28]. In contrast, *M. longipes* females and both sexes of all other species show scaling relationships between leg and body length that are isometric or near isometry (Figure 1B; Supplementary table 1). Taking the two traits individually, *M. longipes* male legs are both significantly longer and more variable than female legs (Figure 2A, B; Supplementary table 1). In contrast, *M. longipes* body size is significantly more variable in males than in females, but average body length is not significantly different between the sexes (Figure 2A, C; Supplementary table 1). Despite these major differences, both sexes presented leg and body length distributions that were not significantly different from normality (Supplementary table 2).

**Figure 1:**
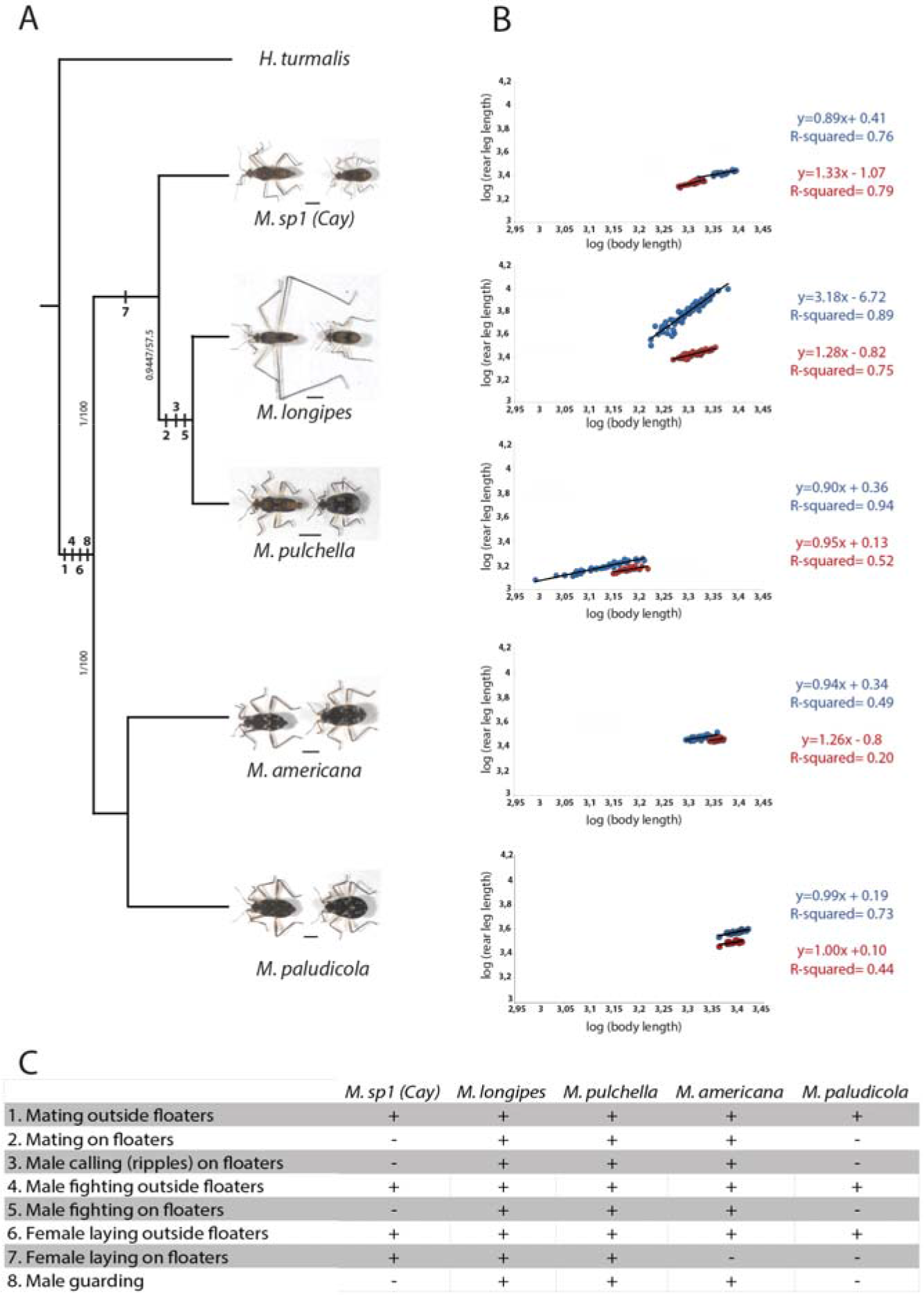
Diversity in leg sexual dimorphism and mating behaviours in *Microvelia*. A) Phylogenetic relationships between five *Microvelia* species using Maximum Likelihood and Bayesian analyses. Support values obtained after Bayesian posterior probabilities and 1000 bootstrap replicates, respectively, are shown for all branches. Pictures of males (right) and females (left) illustrate divergence in sexual dimorphism in the five *Microvelia* species. Scale bar represents 1mm. B) Scaling relationships of log-transformed data between rear legs and body lengths were estimated in males (blue) and females (red) of the five *Microvelia* species using Major Axis regressions. The equations and fitting (R-squared) of the linear regressions in males and females were indicated using the same colour codes. C) Behavioural characters describing the mating system of the five *Microvelia* species. These characters were mapped onto the phylogeny based upon the parsimony criterion.

**Figure 2:**
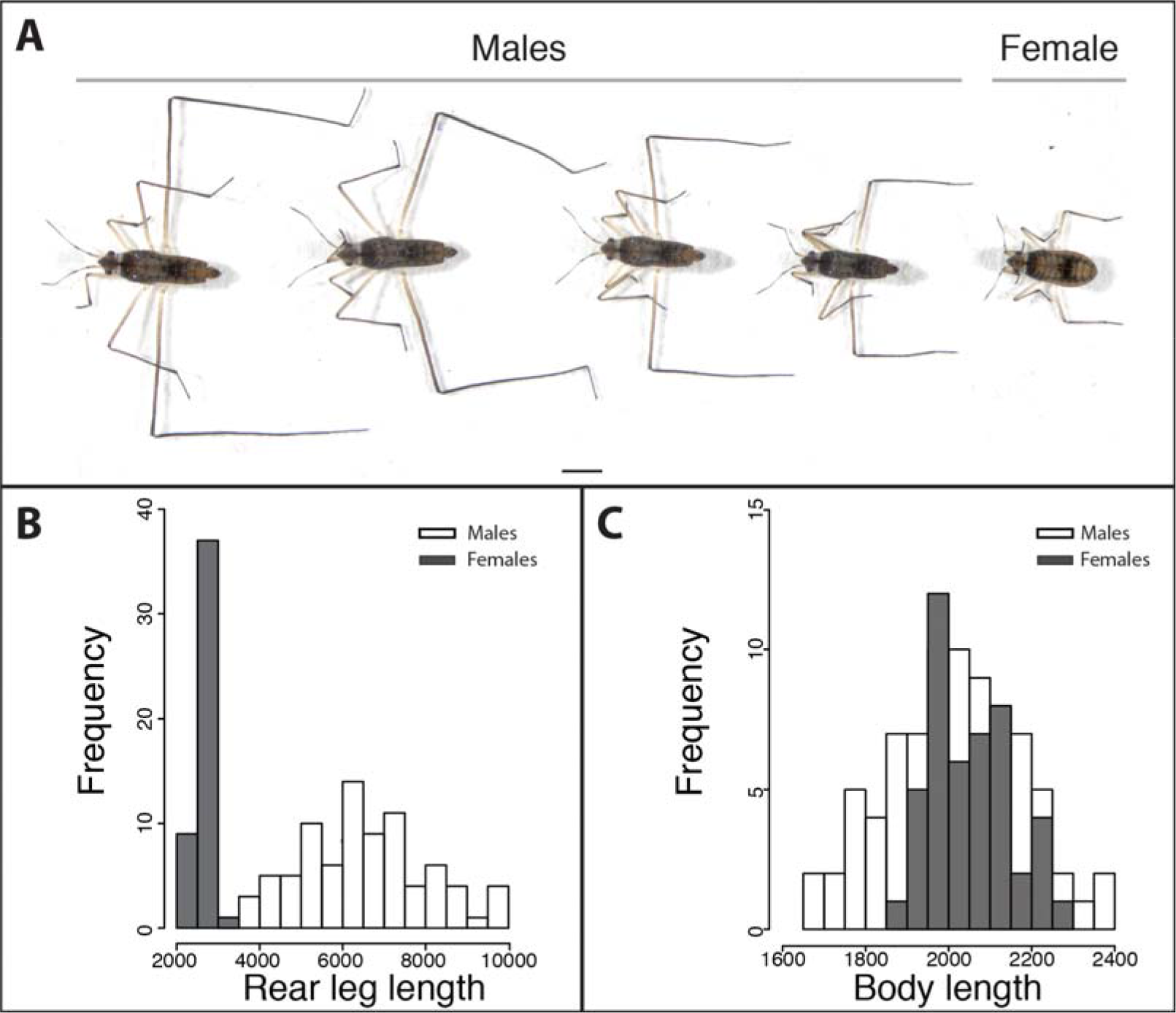
Phenotypic variation of rear leg exaggeration and body length in *M. longipes*. A) Phenotypic variation of rear leg length in males and in a female. (B) Rear leg length and (C) body length distributions of males (white) and females (grey) from a natural population collected in French Guyana.

Finally, we found that the males of three *Microvelia* species (*Microvelia sp.*, *M. americana* and *M. paludicola*) evolved prominent spikes on the rear legs indicative of a function in grasping females during pre-mating struggles [29] (Figure 1A). Overall, these analyses indicate that the evolution of hypervariable exaggerated legs in *M. longipes* males results from the high variance in body length and the associated hyperallometric relationship with leg length (Figure 1 and 2). In *M. pulchella*, despite the high variation in male body length, the near isometric relationship between leg and body length makes their legs less exaggerated and less variable than *M. longipes* males (Figure 1B; Supplementary table 1). Moreover, the diversity in sexual dimorphism between *Microvelia* species does not seem to follow any particular phylogenetic pattern (Figure 1), suggesting that variation in the ecology, behavior, or mating systems may play a role in the divergence of the sexes in these species.

### Mating systems in *Microvelia* species

We characterized mating systems and sexual interactions in all five species to better understand the differences in sexual dimorphism (Supplementary figure 2). In nature, the *Microvelia* genus comprises species that occupy a wide variety of habitats [27], Most species live nearshore, in stagnant, large water bodies [27], Some species, like *M. longipes*, *M. pulchella* or *Microvelia sp*. are gregarious and specialize in small temporary puddles filled with rainwater in tropical South America [27, 30]. Behavioral observations both in the wild and in laboratory-recreated puddles revealed that *M. longipes* males are highly territorial and tend to aggressively guard floating objects consisting of small twigs or pieces of dead leaves (Supplementary figure 3). These are egg-laying sites where males signal to attract females, by vibrating their rear-legs and pounding with their genitalia on the water surface producing ripples (Supplementary videos 1 and 2). We hereafter refer to these objects as egg-laying floaters. When a female approaches the floater, the dominating male switches from signaling to a courtship behavior. The female inspects the floater and either leaves or mates without any resistance with the courting male and immediately lays 1 to 4 eggs (n=26 mating events) (Supplementary figure 3; Supplementary video 2). The male then initiates an aggressive guarding behavior by turning around the egg-laying female and chasing other approaching males to (Supplementary video 2). After egg-laying the female leaves and the male initiates another cycle of signaling on the same floater. During this entire process, other males constantly challenge the signaling male in an attempt to dominate the floater. During these contests, the dominant and the challenging male fight back-to-back by kicking each other with their rear-legs until one of them is chased away (Supplementary video 2). We also observed that females could lay eggs in the mud at the margin of the puddle and that males attempt to mate outside floaters by jumping on female’s back randomly in the puddle.

*M. pulchella*, the sister species of *M. longipes* (Figure 1A), is also found in small temporary puddles [30] and displays a highly similar mating behavior despite the lack of rear-leg exaggeration (Figure 1C). Males of *M. pulchella* compete for egg-laying floaters, fight with their rear-legs, and generate ripples to attract females. Also like *M. longipes*, females of *M. pulchella* lay their eggs both on floaters and in the mud. In spite of similarities in their mating behavior, these two sister species display significant morphological differences, raising the question as to which factors drove the evolution of trait exaggeration in *M. longipes*.

In the three other species, *M. americana*, *M. paludicola*, and *Microvelia sp.*, males possess grasping spines on their rear-leg femurs (Figure 1A) and actively harass females in an attempt to mate. Females consistently struggle through vigorous shaking, frequently resulting in the rejection of the male. Males of these three species also fight occasionally but the fights do not seem to result in the dominance of any particular localized resource (Figure 1C; Supplementary figure 2). *M. americana* and *M. paludicola* females lay eggs exclusively on water margins while females in *Microvelia sp.* lay eggs randomly either on floaters or water margins, but do not do so immediately after mating (Figure 1C; Supplementary figure 2). Altogether, these data show that the behavior, consisting of contests using the rear-legs, predates the origin of exaggerated leg length and could therefore be necessary but not sufficient for its evolution. Moreover, differences in egg-laying habits may have driven the diversity in male mating strategies and sexual dimorphism in the *Microvelia* genus. In small temporary habitats, eggs laid in the mud are at high risk of desiccation when water levels go down, and nymphs tend to drown at hatching when water levels go up, something we frequently observe in laboratory conditions. Egg-laying behavior on floating objects, which remain on the surface despite fluctuating water levels, is likely an adaptation to the fast-changing state of the habitat. Interestingly, male behavior consisting of dominating these egg-laying floaters is observed only in species where females lay eggs just after mating, indicative of the high fitness value in accessing them. This behavior is also associated with the high body length variation in *M. longipes* and *M. pulchella* males (Figure 1B), suggesting a link between body size variation and competition for oviposition sites.

### Intensity of male competition in *M. longipes* compared to *M. pulchella*

In order to evaluate the contribution of exaggerated leg length to male mating success, we tested whether a correlation existed between male leg length and their ability to dominate egg-laying sites. We found increased rear leg length to be strongly correlated with the fighting outcome (ANOVA, F(1, 13) = 144.6, p < 0.01), where the males with longer legs won 97% of the fights (n= 75 fights) and dominated the floater (Figure 3A). We also observed this male dominance over egg-laying sites in *M. pulchella*, which did not evolve leg exaggeration. We therefore hypothesized that male phenotypic differences between *M. longipes* and *M. pulchella* could be driven by differences in the intensity of male competition. When we measured the intensity of male competition in standardized space conditions, we found that *M. longipes* males fought on average 8 times more frequently than *M. pulchella* males in a period of 1 hour (Figure 3B, Supplementary table 3). This indicates that male competition is significantly higher in *M. longipes* than in *M. pulchella.* More importantly, 81 % of *M. longipes* fights occurred on the floaters (Figure 3B, Supplementary table 3) whereas *M. pulchella* males’ fights occurred randomly on and away from floaters (Figure 3B, Supplementary table 3). The same result was reached when we repeated this experiment in standardized density conditions taking into account size differences between the two species (Supplementary table 3). These data demonstrate first that increased rear leg length in *M. longipes* males favors male dominance over egg-laying sites to better intercept gravid females. While both *M. longipes* and *M. pulchella* males intercept females and compete on those egg-laying sites, competition intensity for egg-laying sites is almost an order of magnitude higher in *M. longipes.* A primary difference between the ecology of these two species is that *M. longipes* specializes in rainwater-filled small puddles while *M. pulchella* is a generalist that can be found in both temporary and more stable water bodies ([31, 32] and personal field observations). This difference in niche specialization has two major impacts on *M. longipes* population structure. First, *M. longipes* populations can reach very high densities confined in a small space, something we observed frequently in the wild and which is not the case for *M. pulchella*. Second, because the water level in the puddle can change rapidly (Supplementary figure 4), floaters represent the safest substrate in terms of survival of the progeny. This may explain why females bounce the floater up and down before they copulate and lay eggs (Supplementary video 2), and why *M. longipes* males are so aggressive in dominating these floaters. The situation is different for *M. pulchella* due to the higher stability of the habitat, making floaters less critical and the survival of eggs in the mud more likely. These ecological conditions favoring high-density populations and floating objects as the more suitable egg-laying substrate may have at least contributed to the high competitiveness observed in *M. longipes*, and thus acted as a driving force for the evolution of the exaggerated leg length for use as a weapon. Both empirical and theoretical models suggest that population density can influence aggressiveness and the intensity of sexual selection [33], and our data show how increased competitiveness can drive secondary sexual traits to reach dramatic levels of expression.

**Figure 3:**
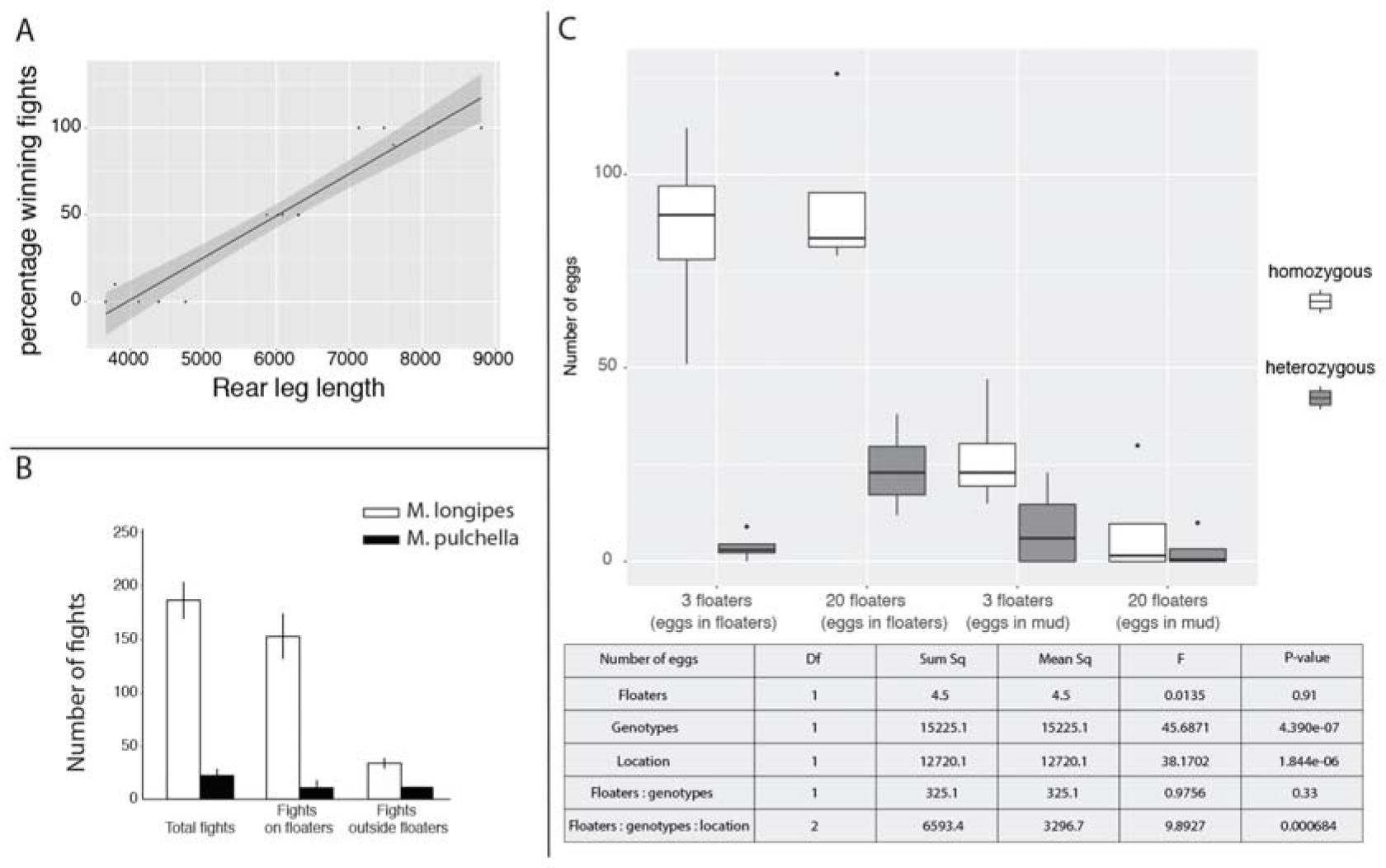
Selective pressures and reproductive fitness of leg exaggeration in *M. longipes* males. A) Relationships between fighting outcome and male rear leg length. Winners correspond to males keeping the access to the egg-laying sites after the fights. Solid line represents a fitted linear regression model, B) Frequency of fights between *M. longipes* and *M. pulchella* on both floaters and outside floaters, C) Fertlization success of large and small males and the contribution of egg-laying sites. Heterozygous eggs result from the siring of short-legged males (short-legs selected line) and females (long-legs selected line). Homozygous eggs result from the siring of long-legged males (long-legs selected line) and females (long-legs selected line). The table summarizes an ANOVA test recapitulating the influence of the floaters and the egg-laying locations on the egg genotypes.

### Effect of exaggerated leg length on male reproductive fitness in *M. longipes*

Post-mating competition is widespread in insects [34], including water striders [35-38], and can strongly alter the outcome of pre-mating strategies [34, 39]. Field observations also indicate that the state of the habitat occupied by *M. longipes* can fluctuate rapidly and, sometimes, the water can evaporate entirely in days (Supplementary figure 4). Moreover, the amount of egg-laying resources is highly variable from one puddle to another and can additionally fluctuate with water level (personal observations from the field). We hypothesized that these rapidly changing conditions will influence competition and mating success across the distribution of male phenotypes. To test this hypothesis, we conducted paternity tests using *M. longipes* lines that are homozygous for distinct microsatellite markers that can reveal the identity of the parents (see methods for more details). We set the experiment such that heterozygous progeny could only originate from eggs fertilized by small males. Because egg-laying floaters represent the primary resource that males dominate to intercept gravid females, we designed a first treatment where floaters were limiting (3 floater for 6 large and 6 small males) and another treatment where floaters were abundant (20 floaters for 6 large and 6 small males). We also genotyped the progeny from eggs laid in the mud to determine mating success of different male phenotypes in contexts other than the dominance of floaters. In all replicates of each treatment, females laid the majority of their eggs on floaters regardless of whether floaters are limiting (91% of a total of 512 eggs) or abundant (71% of a total of 500 eggs) (Figure 3D). However, females laid on average three times more eggs on the mud when floaters were limiting (Supplementary table 4). In the condition where floaters were limiting, small males fertilized 4.6% (15 eggs of a total of 357 eggs) of the eggs laid on floaters and 25% of the eggs laid in the mud (35 eggs of a total of 143 eggs) on average (Figure 3D; Supplementary table 4). Interestingly, the number of eggs sired by small males was more than twice higher in the mud than on floaters (Figure 3D; Supplementary table 4). This suggests that when the dominance of floaters by small males is limited, they primarily achieve egg fertilization by mating outside floaters. In the condition of abundant floaters, the proportion of eggs fertilized by small males on floaters increased significantly to 19% (96 eggs of a total of 468 eggs) (Figure 3D; Supplementary table 4), while that outside floaters remained unchanged (11 eggs of a total of 44 eggs) (Figure 3D; Supplementary table 4). In contrast to the treatment with limiting floaters, here the number of eggs fertilized by small males is almost nine times higher on floaters than in the mud (Figure 3D; Supplementary table 4). These results show that small males can sire significantly more progeny when egg-laying sites are abundant but can also mate outside these egg-laying sites when floaters are limiting. Therefore, sexual selection is strong in favor of large males with long legs but can become relaxed in conditions where egg-laying sites are abundant. Rapid changes in water level and high heterogeneity between puddles are intrinsic to the life history of this species and are expected to cause variation in the amount of accessible egg-laying floaters over time and space. This fluctuating selection is therefore likely to influence the strength of competition and mating success and contribute to the high phenotypic variation found in *M. longipes* natural populations.

### Environmental and genetic contributions to male rear leg variation

We have shown that possible fluctuation in the strength of sexual selection may favor phenotypic variation, however its impact on the mechanistic underpinnings of phenotypic variation in *M. longipes* males is unknown. We therefore tested how genetic variation and phenotypic plasticity contribute to the maintenance of high variation in *M. longipes* male leg length. Artificially selected large and small male lines, generated through 15 sib-sib successive crosses from a natural population, showed a shifted distribution of male leg length towards the respective extreme phenotypes of the distribution (Figure 4A). The difference between these two lines held for both absolute and relative leg length, but the allometric coefficient remained, nonetheless, unchanged (Supplementary figure 5; Supplementary table 5). This shows that genotypic variation contributes to the variation in both rear leg length and body size.

**Figure 4:**
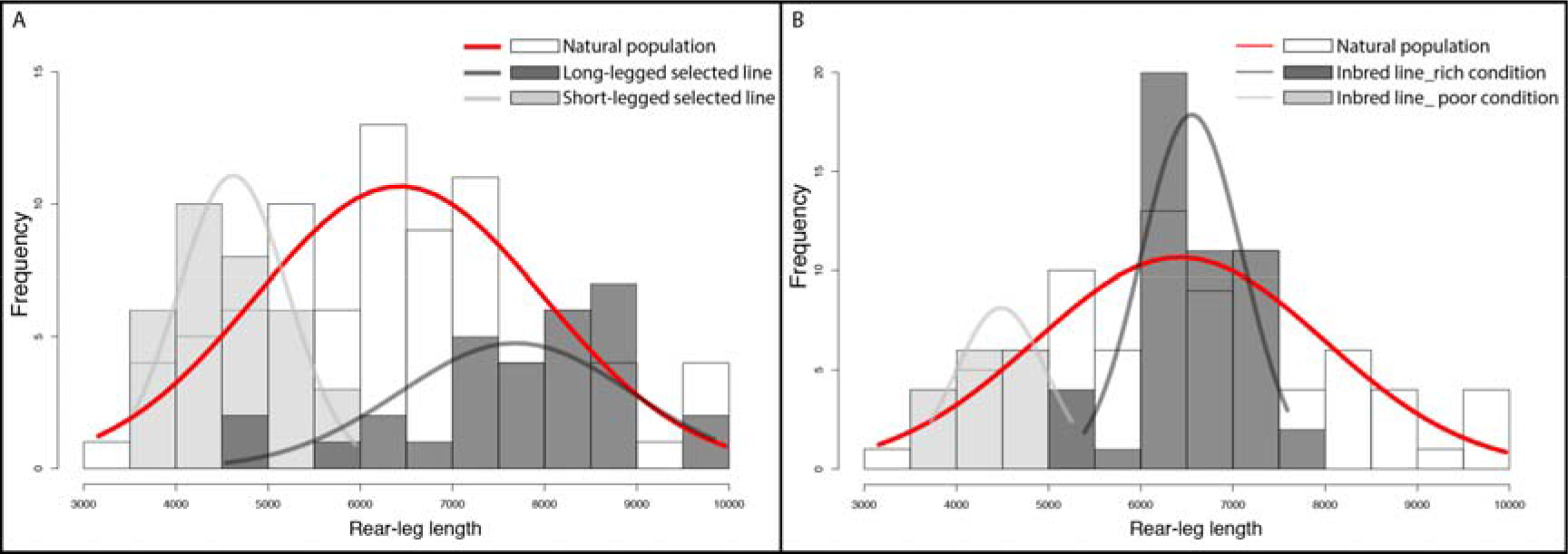
Environmental and genetic contributions to rear leg length variation in males *M. longipes.* A) Rear leg length distributions of adult males from natural population (white) and from an inbred line that developed under poor (light grey) and rich (dark grey) conditions. B) Rear leg length distributions of adult males from natural population (white) and from two inbred lines that were selected for short (light grey) or long (dark grey) rear legs under rich condition. Normal curves were fitted to each distribution after testing for normality of each condition.

Next, we tested the reaction norm of one of these inbred lines in poor and rich nutritional condition. Despite near identical genotype, individuals reared in poor condition developed shorter legs than individuals reared in rich condition such that the distributions of the two treatments were almost non-overlapping (Figure 4B, Supplementary figure 6; Supplementary table 5). Importantly, this difference in leg length between the two treatments resulted mostly from differences in overall body size (t-test body length: t = 10.5643, df = 25.274, p-value = 9.244e-ll) but not in the scaling relationship as we failed to detect any significant difference in the allometric coefficient or the intercept between rich and poor conditions (Supplementary figure 6; Supplementary table 5). The same result was reached when we tested condition dependence in a laboratory population where no specific selection has been applied, although some statistical tests detected a small but significant difference in intercept between the two conditions (Supplementary figure 7; Supplementary table 5). This difference was nonetheless not significant when using a linear model (ANOVA, F(1,88)= 2.6202, p-value= 0.1076). We therefore conclude that, in *M. longipes*, male body size is highly condition-dependent but the rear legs are not or they are to a small extent after body size correction. Altogether, these results suggest that male leg length variation in nature results from the contribution of both genetic variation and strong condition dependence. The fluctuations in the amount of egg-laying floaters, combined with phenotypic plasticity, is expected to result in the maintenance of a certain degree of genetic variation in the population through the incomplete removal of alleles of small leg and body size. However, episodes of relaxed selection are not only known to increase genetic variation in the population, but also to favor the evolution of reaction norms and therefore increase phenotypic plasticity [40, 41].

## Conclusions

This study provides a good example of how various ecological factors influence the intensity of sexual selection and ultimately the mechanisms and patterns of phenotypic variation. In the genus *Microvelia*, mating systems are diverse and are likely to influence the diversification of male-specific secondary sexual traits used in pre-mating copulatory strategies. The intense male competition to dominate egg-laying sites in *M. longipes*, unlike other *Microvelia* species, underlies the evolution of exaggerated leg length used as a weapon. Dominating males that intercept and copulate with gravid females on egg-laying sites gain a significant increase in their reproductive fitness by siring the majority of the eggs. This intense selection on increased leg length can, however, be relaxed when egg-laying sites are abundant thus allowing small males to fertilize a significant number of eggs. We have also shown that plasticity in response to nutritional condition along with genetic variation both contribute to the high phenotypic variation we observe in body and leg length. It is possible that fluctuating selection, combined with phenotypic plasticity, both facilitate the dramatic increase and maintenance of phenotypic variation in *M. longipes* compared to other *Microvelia* species. It is also important to note that the fluctuating selection described here (availability of egg-laying floaters) is independent of the individual condition. Therefore its influence on phenotypic variation cannot be the consequence of a pre-existing increase of condition-dependence, as it would be the case for fluctuating selection on food resources for example. Altogether, these results point to two ways in which alleles for small male body and leg size will be maintained in the population. First, because small males can sire a significant number of progeny due to possible episodes of relaxed selection. Second, because males with allelic combinations for low trait expression can develop larger body and leg size if they experience higher nutritional condition during development. Therefore condition dependence causes a non-linear relationship between genotypes and phenotypes, making directional selection less efficient in depleting genetic variation. In their opinion paper, Cornwallis and Uller [23] refer to this process as a “feedback loop between heterogeneity, selection and phenotypic plasticity”.

The findings outlined here open important research avenues to gain a general understanding of how sexual selection can impact phenotypic evolution. *Microvelia longipes* as a new hemimetabolous insect model with an exaggerated secondary sexual trait offers the opportunity to complete the substantial literature in holometabolous insects such as beetles or various flies [8, 42–45]. Males of many species of water striders employ water surface ripples as mating calls, and it is unknown whether females can deduce the size of the male from the ripple pattern and whether this would influence female choice [27, 46–49]. In addition, the number and the frequency of allelic variants underlying this trait and how they may interact with the environment remains to be tested. The ease of rearing and the relative short generation time make *Microvelia longipes* a powerful future model to study the extent to which genetic variation and environmental stimuli influence gene expression and ultimately phenotypic variation.

## Acknowledgements

We thank Emília Santos and Antonin Crumiére for help with collecting bugs, Felipe Moreira for help with species identification, Russell Bondurianski, Locke Rowe, Kevin Parsons, Gaël Yvert, François Leulier, Augustin Le Bouquin, Amélie Decaras, Cédric Finet, Aidamalia Vargas, Roberto Arbore for helpful discussions and comments on the manuscript, and David Armisén and Antoine Melet for help with genetic markers identification and data measurements. This work was supported by an ERC-CoG# 616346 and labex CEBA to AK, and a PhD fellowship from Ecole Doctorale BMIC de Lyon to W.T.

## Author contributions

A.K. and W.T. designed research; W.T. performed research; A.K. and W.T. analyzed data; and A.K. and W.T. wrote the paper

## Material & methods

### Population sampling and culture

*Microvelia* populations were collected during fieldwork in French Guyana in Crique Patate near Cayenne. These populations were maintained at 25°C and 50% humidity. The bugs were fed on either frozen or freshly euthanized crickets. Adults laid eggs on Styrofoam floaters and the hatched nymphs were raised in separate tanks to avoid cannibalism.

### Measurement of Microvelia species and statistics

Rear leg and body lengths of all *Microvelia* species were measured with a SteREO Discovery V12 (Zeiss) using the Zen software. All statistical analyses were performed in RStudio 0. 99.486. Comparisons for mean trait size and trait distributions were performed on raw data whereas log-transformed data were used for scaling relationship comparisons. We used Major-Axis (MA) regression to assess differences in scaling relationships (“smatr” package in R, [50]). Differences in intercepts were estimated using a Wald statistic test and we used Likelihood ratio test for differences in slopes [50].

### Behavioral observations and video acquisition

Male and female interactions of all *Microvelia* species were observed in a recreated small puddle, using local mud, and were filmed with a Nikon digital camera D7200 with an AF-S micro nikkor 105mm lens. Observations and video acquisitions were taken a couple of hours after the bugs were transferred to the puddle. In *M. longipes* and *M. pulchella* male and female interactions were also observed in the field.

### Microvelia phylogenetic reconstruction

The phylogenetic relationships between the five *Microvelia* species used in the behavioral assays was generated using the Geneious software version 7.1.9 using plugins MrBayes version 3.2.6 and PhyML version 3.0, as described in [51]. The phylogenetic reconstruction was performed using 14 molecular markers retrieved from in house transcriptome databases: *12S RNA*; *16S RNA*; *18S RNA*; *28S RNA*; *Cytochrome Oxydase subunit I (COI)*; *Cytochrome Oxydase subunit II (COII)*; *Cytochrocme Oxydase subunit III (COIII)*; *Cytochrome b (cyt b)*; *Ultrabithorax (Ubx)*; *Sex combs reduced (Scr)*; *Gamma interferon inducible thiol reductase (gilt)*; *Antennapedia (Antp)*; *Distal-less (dll)*; *Doublesex (dsx)*. All these markers can be retrieved in GenBank using the following accession numbers: (Will be provided before publication). Phylogenetic reconstruction was perfomed using MrBayes version 3.2.6 and PhyML version 3.0 in Geneious 7.1.9 as described in [52], Concatenation of sequence alignments and phylogenetic tree in Newick format are also available in the Dryad Digital Repository.

### Fight frequency assay

To compare the number of fights between males of *M. longipes* and *M. pulchella*, we isolated twenty-five adult males and females over a period of two days. Both sexes were then mixed together in the puddle during 30 minutes before observation. The number of fights on and outside floaters was counted for a period of one hour. We repeated the experiment the following day with the same males and females kept mixed together overnight. Finally, in order to account for population density, because of size differences between the two species, we calculated the number of fights in a reduced sample of ten males and ten females in *M. longipes*.

### Artificial selection experiment, phenotyping and line sequencing

We assessed the genetic contribution of rear leg length variation in males by performing an artificial selection experiment for long versus short-legged males. Individual males from the French Guyana natural population were selected for their absolute rear leg sizes and mated with random females to initiate the successive sib-sib crosses. After fifteen generations of sib-sib inbreeding, two populations selected for extreme phenotypes were amplified over two generations before phenotyping.

### Condition-dependence experiment

First instar nymphs were collected just after hatching and individuals were reared attributed in either poor or rich nutritional condition. In the poor condition, hundred first instar nymphs of the long-legged inbred line were fed everyday with ten crickets during the first two nymphal instars, followed by only three cricket legs until adulthood. In the rich condition fifty individuals of the same line were fed with ten crickets, changed everyday, over their entire nymphal development until adulthood. In a second experiment we tested the effect of condition in an independent set of individuals from the lab population. This experiment was performed on three replicates per condition, with fifty individuals per condition. Replicates were then pooled for the analysis. We started the poor condition by feeding the first two nymphal instars with height crickets everyday and then switched to one small cricket every two days until they reached adulthood. Individuals from the rich condition were fed during their entire nymphal development with eight crickets everyday.

### Microsatellite development

DNA from *M. longipes* was extracted from ten male and female individuals from the lab population. Insects were first frozen in liquid nitrogen before DNA extraction with the Genomic DNA Buffer Set kit from Qiagen. We used 12[ig of DNA for sequencing on an lon-Torrent Sequencer machine (Sequencing Plateform IGFL, Lyon, France) generating 3.7M reads with median size of 317 bp.

We used the program Exact Tandem Repeat Analyzer 1.0 (available from ftp://ftp.akdeniz.edu.tr/) in order to identify reads containing microsatellite repeats [53]. The software also provided primers for microsatellite amplification (Supplementary table 6). Forty-seven markers of various tandem repeats were tested by PCR and forty of them were successfully amplified from an aliquot of the genomic extraction used for the Ion-Torrent sequencing (Supplementary table 7). We also tested the same pairs of primers on DNA extraction from single individuals of the stock population (Supplementary table 7), as described in [54].

In order to identify polymorphic microsatellites, we first performed PCRs of these forty markers on fifteen individuals from the lab population and sent these samples for genotyping (Genoscreen, Lille, France). We considered polymorphic, any marker that was showing at least two genotypes in at least two individuals. Nine polymorphic reliable markers could be identified, among which one of them (microsatellite 21, see Supplementary table 6) was showing a polymorphism between our two lines of interest. This microsatellite was genotyped in seven males and seven females from each line to confirm the polymorphism prior to the paternity test experiment.

### Paternity test

To assess the fertilization success of long and short-legged males, we collected six males from both the short- and the long-legged inbred lines, and put them together in an artificial puddle with twelve females from the long-legged inbred line. We conducted two treatments, each with four replicates, where we provided twenty or three floaters in the puddle to create conditions with abundant and limiting egg-laying sites, respectively. On day 3, the parents were collected, their DNA extracted and the microsatellite of interest amplified for genotyping. We then isolated the floaters and genotyped the nymphs that hatched from the floaters and those that hatched from the mud after adults and floaters were removed.

---

**Supplementary figure 1:** Comparative morphology of the legs across *Microvelia* species sample in figure 1. Note the presence of grasping traits on male legs in *Microvelia americana, Microvelia paludicola* and *Microvelia sp.* These traits are absent in *Microvelia longipes* and *Microvelia pulchella*.

**Supplementary figure 2:** Schematic summary representation of the mating systems in the five *Microvelia* species

**Supplementary figure 3:** *M. longipes* natural habitat. Top panel: Example of rain-filled puddle in French Guyana in Crique Patate near Cayenne where *M. longipes* population was collected. Middle panel: Zoom on the floating substrates deposited on the water surface of the puddle. Bottom panel: Example of floater full of *M. longipes* eggs. Scale bar represents 5mm.

**Supplementary figure 4:** Fluctuating environment. Pictures of a rain-filled puddle, over a five-day period, in Rio de Janeiro where a *M. longipes* natural population was found.

**Supplementary figure 5:** Artificial selection effects on body-leg scaling relationships. Static allometry on log-transformed data between rear leg and body lengths for adult individuals selected for long (black) and short-legged (grey) males.

**Supplementary figure 6:** Nutrition effects on body-leg scaling relationships in the longlegged inbred line. Static allometry on log-transformed data between rear leg and body lengths for inbred adult individuals fed on rich (black) and poor (grey) diets.

**Supplementary figure 7:** Nutrition effects on body-leg scaling relationships in the lab population. Static allometry on log-transformed data between rear leg and body lengths for unselected adult individuals fed on rich (black) and poor (grey) diets.
**Supplementary table 1:** Morphometric data and associated statistical tests. Summary table of the adult measurements and statistical tests for all *Microvelia* species.

**Supplementary table 2:** normality tests: Tests for normal distribution in all *M. longipes* conditions. Values of Shapiro tests and associated p-values for each *M. longipes* adult population reared in different condition.

**Supplementary table 3:** Fight frequency between *M. longipes* and *M. pulchella* males. Summary table of the number of fights in *M. longipes* and *M. pulchella* males in different conditions for a period of one hour. Below are the associated statistical tests for differences in fight frequency.

**Supplementary table 4:** Count and genotype of the total number of eggs laid by females in each condition.

**Supplementary table 5:** Statistical tests associated with differences in leg length and scaling relationships between artificially selected and nutritionally manipulated *M. longipes* populations.

**Supplementary table 6:** Table containing the primer sequences, the full sequence, the motif and the length of each tested microsatellite.

**Supplementary table 7:** Table PCR protocols for microsatellite amplifications and single individual genotyping.

**Supplementary video 1:** *M. longipes* male vibrations in slow motion.

**Supplementary video 2:** *M. longipes* mating system.

